# Unpredictable Prey Motion Shapes Behavior Across Timescales

**DOI:** 10.64898/2026.05.18.726036

**Authors:** Sean C. Cadigan, Nathon A. Smith, Te K. Jones, Melville J. Wohlgemuth

**Affiliations:** University of Arizona; Johns Hopkins University

**Keywords:** echolocation, tracking, behavioral, bat, computational, auditory

## Abstract

Locating, tracking, and intercepting objects is a fundamental behavior for many organisms. For instance, predators must track and capture erratically moving prey for their survival. Using the echolocating bat as a model species, we investigate how short-term changes in target motion predictability affect longer-term motor plans when tracking a prey item. We used a paradigm where prey motion is under experimental control, and then applied computational methods to characterize how target motion predictability influences short- and long-term behavioral control. We find that target motion predictability during the tracking phase of insect capture influences both short-term changes in sonar call control, as well as longer-term behavioral control for transitioning between hunting phases. For changes in immediate behavioral control, bats produce more bursts of calls at a higher rate when tracking unpredictable moving prey, an indication that the bat is collecting more information about the target’s motion for unpredictable than predictable trials. In terms of longer-term behavioral control, target motion unpredictability delays the transition from tracking to capture phase behaviors. We suggest that the bat does this to collect more information about target motion to time the transition from tracking to capture behaviors for hunting success. Additionally, we find the effects of target motion unpredictability are first seen as changes in the vocal motor plan and then the auditory motor plan (ear motion), hinting at a sequencing of motor changes that warrant further investigation.

**Summary:** When presented with a more challenging hunting task, bats will increase their production of bursts of calls at a higher rate and delay their transition into capture behaviors.

## Introduction

Success in tracking and intercepting a moving target is improved by updating the target’s relative location over time to appropriately control behaviors for capture (Thyselius et al., 2023). Many animals, including invertebrates like dragonflies (Talley et al., 2023), gather information over time to predict the future location of a target. However, this process is more complicated if the target is moving unpredictably, and more sensory information may be needed to properly track and capture an erratically moving object. A common example is trying to catch a frisbee thrown in windy conditions - the trajectory is difficult to predict, and more information is needed for interception. Predators often face similar challenges when tracking and capturing evasive prey, or under changing sensory conditions. Past work on the archerfish shows how it adapts moment-by-moment behaviors based upon changing sensory conditions (Volotsky et al., 2024), but how sensory uncertainty affects longer-term motor planning is less understood. The current study utilizes the hunting patterns of the echolocating bat as a model for how behavior is adapted across short and longer timeframes to track objects moving unpredictably. We hypothesize that moment-by-moment calculations about target motion ultimately influence both short-term motor control to increase sensory acquisition, as well as longer timescale planning for the goal of intercepting a target. For this study, we define short-term changes in motor control as the integration of a single echo’s auditory information into changes in the subsequent sonar call and motion of the ears. In terms of longer-term behavioral adaptations, we define these as the dramatic behavioral adaptations that are made as the bat transitions from one hunting mode to the next. These hunting phases are based off of prior work (Moss and Surlykke, 2010, see Figure 1A). For our study, we focus on longer-term changes in behavior related to the transition from tracking behaviors to capture behaviors. We therefore predict that when animals track erratically moving objects, they will alter long-term behavioral control to delay the initiation of capture behaviors to collect more information about object motion. We will test this hypothesis using echolocating bats as they adapt their sonar imaging behaviors to track targets moving predictably and unpredictably.

**Figure 1.**
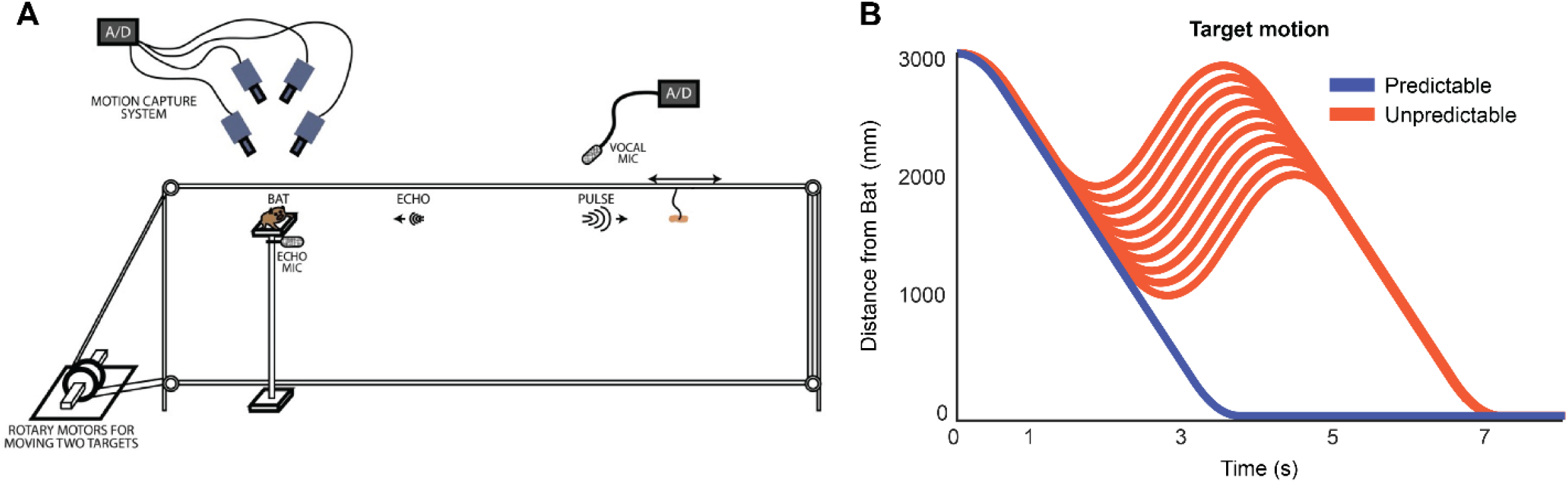
A) Platform set-up with motion tracking cameras above, four pullies linked to a motor, and a microphone to record vocalizations and echoes. B) Target distance from the bat as a function of time. The bottom blue line is referred to as predictable, others are unpredictable (orange).

On a fundamental level, the bat produces ultrasonic calls that reflect off of objects in the environment and return echoes to the bat’s ears. These echoes are used to identify the target, and then coordinate hunting behaviors. As the bat first searches, then tracks, and finally captures prey, its active sensing behaviors are optimized to extract task-relevant information (Jakobsen et al., 2013; Kothari et al., 2018; Macías et al., 2016). In terms of active sensing for echolocation, the bat produces and adapts the signal for probing the world (i.e., sonar calls, Moss & Surlykke, 2010), as well as the positioning of its external auditory system when receiving echoes (Griffin, DR. Dunning, DC. Cahlander, DA. Webster, 1962; Pye, J D. Roberts, 1970; Wohlgemuth et al., 2016). One adaptation made to sonar calls is a change in the rate of production. For example, when hunting, the bat increases call rate with decreasing target distance to increase the rate of echo returns (D. Griffin, 1958). For both insectivorous and frugivorous bats, times of increased sensory uncertainty often correlate to bursts of calls at a higher rate, termed sonar sound groups, or SSGs (Hulgard & Ratcliffe, 2016; Kothari et al., 2014, Beetz et al, 2019). These adaptations increase the number of sensory snapshots (echoes), and ultimately the amount of information about the environment, as motor demands increase. In this way, when bats experience sensory uncertainty, they produce more calls to increase the rate of echo returns.

In addition to changes in call rate with decreasing target distance, prior work has shown how the separation between the outer ears of the bat (i.e., inter-pinna separation) increases as the target distance decreases (Wohlgemuth et al., 2016). Modeling work has demonstrated that a narrow inter-pinna position focuses auditory attention narrowly in front of the animal, while a wider separation broadens auditory sensitivity to more peripheral locations (Gao et al., 2011). Adaptations of sonar call timing and ear positioning allow the bat to modify outgoing sensory signals and echo reception, respectively, for target tracking and capture (Moss et al., 2011; Walker et al., 1998). Past work has examined sonar call design as bats tracked targets moving in range (Aytekin et al., 2010; Hartley, 1992; Moss & Schnitzler, 1995a), but how target motion predictability influences sonar call design and echo reception has not been examined. The current study characterized the different adaptations in sonar call timing and inter-pinna separation when tracking predictable and unpredictable moving targets.

The bat transitions through several discrete behavioral modes while hunting, with small changes in behavior within a mode, and dramatic shifts in behavior between modes (Geberl et al., 2015; D. R. Griffin et al., 1960). The hunting modes of interest for our investigation include the tracking and capture phases, and how target motion predictability affects the timing of this transition. As the bat prepares for capture, it dramatically and abruptly increases the rate of call production to almost 200 calls per second (D. Griffin, 1958). There is a similar abrupt change in the positions of the outer ears immediately before capture (Griffin et. al., 1962). Performing this transition from tracking to capture mode at the appropriate target range is critical. The bat must transition from tracking to capture phase at the appropriate time; if the bat shifts from tracking to capture too early or too late, it will miss the target (Geberl et al., 2015; D. Griffin, 1958). Some organisms exhibit a fixed action pattern when hunting, like the toad, where the prey stimulus initiates a stereotyped sequence of capture behaviors, even if the prey item is removed the last second (Ewert, 1970). We suggest that the bat actively controls the transitions of its hunting behaviors, specifically the tracking and capture phases. We therefore hypothesize that if a target’s motion is predictable, the bat will transition from tracking to capture phase at further target distances because it has more reliable location information. However, if target motion is unpredictable, we hypothesize that the bat will wait to collect more information about the target’s position before transitioning to capture behaviors, and therefore perform this transition when the target is at closer distances. This hypothesis was tested using a paradigm where the target’s motion was controlled by a motor that interfaced with a computer. The motion of the motor could be controlled by the experimenter, thus allowing motion predictability to be experimentally manipulated. Through this approach, we find that bats wait to shift from target tracking to capture behaviors when the prey is moving unpredictably (i.e., shift to capture at closer target distances for unpredictable motion). This finding lends support to the idea that more sensory information about the target’s position is needed when its motion cannot be predicted, leading to delays in capture behaviors to collect more sensory information and ensure success.

## Methods

### Behavioral Training

Bats involved with this study were acquired in Maryland with permission from the Department of Natural Resources and housed in a vivarium on the Johns Hopkins University campus. All procedures and experiments were approved by the Johns Hopkins University Institutional Animal Care and Use Committee (IACUC).

Three big brown bats (*Eptesicus fuscus*) were trained to sit on a platform and track a prey target *(Tenebrio molitor* larva) as it moved towards the bat via a stepper motor (Aerotech NEMA 17) driving a monofilament wire (Fig. 1A, also see Wohlgemuth et al., 2016). This wire was wound around four pulleys that provided control of target motion along the range axis from the bat. Motion parameters, such as velocity, acceleration, and timing, could be precisely determined by custom software written in MATLAB.

Bats were initially trained by pairing an auditory stimulus to a food reward placed on the suspended wire immediately in front of the platform. After the association between the stimulus and food reward was learned, a delay between the stimulus and reward was introduced and slowly increased as the starting distance of the prey from the bat was increased. Once the starting position for the prey reached 3.0m, and the bats’ vocal behavior exhibited characteristics of active tracking (e.g., increase in call rate with decreasing target distance), catch trials were included in the trial sequences in approximately 30% of trials. The catch trials entailed hanging the prey at the same start position but not connected to the motor system. The motor system was then activated, but the prey item remained at the start position. By hearing the motor without target motion, the bat unlearned the association of target motion with the sound of the motor. This ensured tracking behavior was a result of target motion and not a learned association with the sounds produced when driving the motor. Once the bats learned the task and were changing their vocal behaviors with changing target distance, we started the predictable/unpredictable trial types and no longer presented catch trials.

### Target Motion

Eleven target motion conditions were used to move the prey along its path toward the bat, and these 11 motions can be categorized into two major target motion varieties: predictable and unpredictable. What is referred to as “predictable” motion delivers the prey item to the bat with no change to the direction of travel over its 3-meter path (Fig 1B, blue line), reaching a peak velocity of 4 m/s, approximately the same speed as a bat foraging on the wing (Hayward & Davis, 1964). Target motion labeled as “unpredictable” is designed to limit the bats’ ability to anticipate the target’s future position. This was done by interleaving ten different target motions where the target unexpectedly reverses the direction of travel when the target was in the range of 1 to 2 meters from the bat (Figure 1B, orange lines). For any given trial, one of the 11 target motion conditions were chosen at random to ensure the bat tracked the prey using its sonar imaging system and had no other predictions of the target’s motion beyond what was sampled with each echo.

### Data Acquisition

Sonar vocalizations and the positions of the pinnae were recorded as the bat performed its target-tracking behaviors. Synchrony across recording devices was achieved via an analog 5-volt square pulse sent from the program controlling the motor to the audio and video systems. This signal initiated the respective recording processes at the start of the target motion and continued for a duration of 10s. A buffer was built into the recording system that ensured at least an additional 2s of recorded behavioral data after the target arrived at the bats’ position.

Vocalizations were recorded by ultrasonic microphones, manufactured by Ultra-Sound Advice, placed above the target path 3.2m from the bat. Auditory data was received by a preamplifier, filtered for frequencies 15 – 110 kHz (Kemo VBF44), and digitized by a National Instruments M-series data acquisition board at a sample rate of 250 kHz. Data was stored on a hard drive for later analysis.

Motion tracking cameras, Vicon Nexus Motion Tracking System, were used to record the 3D positions of the head and ears. Four T-40s series cameras were mounted above the bat that gave optimal field of view overlap of three, 3mm IR tracking dots/markers placed on the tips of the ears and crown of the head. Positions were recorded at a rate of 500 Hz and calibrated to a sub-millimeter accuracy using a moving wand-based calibration method. Each of the cameras records the 2D position of the tracking dots in their field of view and by combining information across cameras using Vicon Nexus software (ver. 1.85), a 3D reconstruction of marker motion can be made.

### Data Analysis

#### Unpredictable motion analysis

The bats were presented with either a predictably moving target, or target motions that were unpredictable. To combine across unpredictable motion conditions, we first had to determine if there were any significant differences between sonar behaviors for tracking each unpredictable motion variety. We analyzed the target distances at which the bats transitioned from tracking to capture behaviors across the 10 unpredictable motion conditions (Figure S1) for both sonar calls and interpinna separation. We then used a 1-way repeated measures ANOVA to test for significant differences in behavior across the unpredictable motion conditions.

#### Audio analysis

Calls were identified within trial data using custom software in MATLAB run by lab personnel. An amplitude threshold was drawn through a low-passed, rectified version of the original oscillogram where the times of call onset and offset were identified. These automatic determinations of call onset and offset were then manually checked for accuracy and modified as needed. Call times were then corrected to account for the travel time of sound in air. The temperature and humidity of the experimental space were recorded each day to ensure accurate calculations of sound velocity. Sonar call interval was calculated by subtracting the times of successive vocalizations and call duration was calculated by subtracting offset and onset times of the same calls. The call rate is calculated by taking the inverse of the call interval. Trials were excluded from further analysis if the bat did not adapt call rate with target distance.

Once the onsets and offsets of each call were determined, several additional analyses were performed on the audio data. Summary plots were generated by averaging call rate data across trials within bins of target distance. From 3100 – 0mm, 100mm-wide bins were used to average call rate data for the calls produced within the corresponding target distance. The standard error of the mean was then calculated for each bin within the same trial type (predictable and unpredictable) and plotted. To measure where in the target’s path the average vocal rate differed between the two motion types, a d-prime analysis was conducted to determine when predictable target call rates significantly differed from unpredictable call rates. For this analysis, d-prime is defined as the difference of the means divided by the square root of the average variance. To test for significance, a 95% confidence interval was calculated by pooling data across motion types, randomly splitting the pooled data into two groups, and calculating the d-prime between these randomly selected groups. This was performed 10,000 times to get a shuffle distribution for confidence interval calculations. To confirm the validity of this analysis, we also performed an ROC analysis on the same data (Figure S2A).

On a trial-by-trial basis, we set a criterion for determining the start of capture phase calls. This was defined as the first call in a sequence of calls reaching a call rate above 100 Hz and sustained for more than three calls. Additionally, we calculated the proportion of calls included in sonar sound groups (SSGs) per total calls for each motion type. SSGs are a series of two or more calls identified by their short call intervals which are preceded and followed by longer call intervals (Kothari et al., 2014). By definition, the intervals before and after the sound group must be 20% longer than the mean of the intervals within the sound group. Intervals within the group must be similar in duration, specifically within a 70% error of the mean interval value. These criteria were used to identify the sonar calls produced in SSGs in the target tracking call sequences recorded from each bat. The total number of calls that were used in SSGs were then divided by the total number of calls produced in that trial. Significance between motion types was then calculated via permutation test.

#### Motion analysis

Utilizing the 3D tracking data calculated by the Vicon Nexus software, the left ear, right ear, and top of the head were labeled in all frames where the markers were visible. Smoothing and cubic-spline interpolation were applied to minimize positional noise in the tracking data and fill gaps smaller than 50 milliseconds. The inter-pinna separation, defined as the 3D distance between the markers on the tip of each ear, was calculated throughout the trial. Summary plots of ear motion were created by averaging over trials. From 3100 – 0mm, 100mm-wide bins were used to average the interpolated and normalized inter-pinna separation within the corresponding target distance. Normalization involved taking the average inter-pinna separation of the first 0.5 seconds of target motion and then subtracting this value from the whole trial. The standard error of the mean was then calculated for each bin within the same trial type (predictable and unpredictable) and plotted. To measure where in the target’s path the normalized inter-pinna separation differed between the two motion types, a d-prime analysis was performed as described for the call rate analysis. We also calculated the echo arrival times from the known distance of the bat to the target at each call onset. Prior work (Wohlgemuth et al, 2016) has shown moment-by-moment deflections in outer pinnae position for each arriving echo, and this analysis was to determine if the same behavior was found in the current study.

Prior work has shown that the separation between the pinnae increases dramatically before insect capture, and we derive a quantitative measure of when this increase in inter-pinna separation begins on each trial. The inter-pinna separation data was analyzed to find the base of the longest sustained, widest pinna separation before target arrival. A trial average of pinna separation was calculated and used as a threshold. Times when the inter-pinna separation rose above this threshold were used in a time-weighted average to determine the significance of the outward motion. The greatest average pinna separation above threshold corresponded to the final outward pinna movement, and the local minimum before this position was taken as the start of the capture phase pinna movement.

#### Statistical analyses

All audio and motion data were normalized to target distance to compare across bats and trial conditions. Time was thus converted into “distance from bat” measurements. To verify the significant differences in tracking behaviors employed across unpredictable and predictable motions, a permutation test was used on both the audio and motion capture data. More specifically, data was randomly shuffled into two groups, where, on each iteration, a mean difference was calculated for each randomized group and compared to the difference in mean of the actual groups. To calculate a p-value, the fractional value of differences in random means greater than the difference in actual means in 10,000 iterations is used. A similar permutation test was employed to determine significance in sonar sound groups per total calls across unpredictable and predictable trials. We tested whether a data set was normally distributed using the Lilliefors Test, and then to test the correlations among predictable and unpredictable data, the MATLAB corrcoef function was used which performs a Pearson correlation coefficient calculation.

## Results

In the current platform-target tracking paradigm, three bats were trained on the task, and data was collected over 24 different sessions from each bat. The average number of trials per day was 12, with a minimum of 5 trials and a maximum of 26 trials. The various target motion varieties were randomly interleaved across trials within a day’s session, totaling 304 predictable motion trials and 557 unpredictable motion trials across all sessions and bats. The bats performed characteristic adjustments to prey interception behaviors typical of hunting on the wing; including adjustments to call rate (i.e., calls per second) and purposeful movements of the pinna based on target distance (Figure 2A, D). In Figure 2A, an example oscillogram of a hunting sequence from a ‘predictable’ motion trial is shown. The call rate (calls per second, Hz) increases in relation to decreasing target distance, with an abrupt increase when transitioning from the tracking to the capture phase. In Figure 2B, the call rate is plotted as a function of target distance for an unpredictable target motion. We found that behavioral control across unpredictable motion conditions was not significantly different (Figure S1), and we therefore combined data across unpredictable motions. For both motion types, there is a clear increase in call rate with decreasing target distance, as would be expected for naturalistic hunting conditions (Figure 2A-C). Just before target arrival, the call rate rises rapidly as the bat anticipates imminent target capture, with this increase in call rate increasing the number of echoes returning to the bat’s ears (Figure 2C).

**Figure 2.**
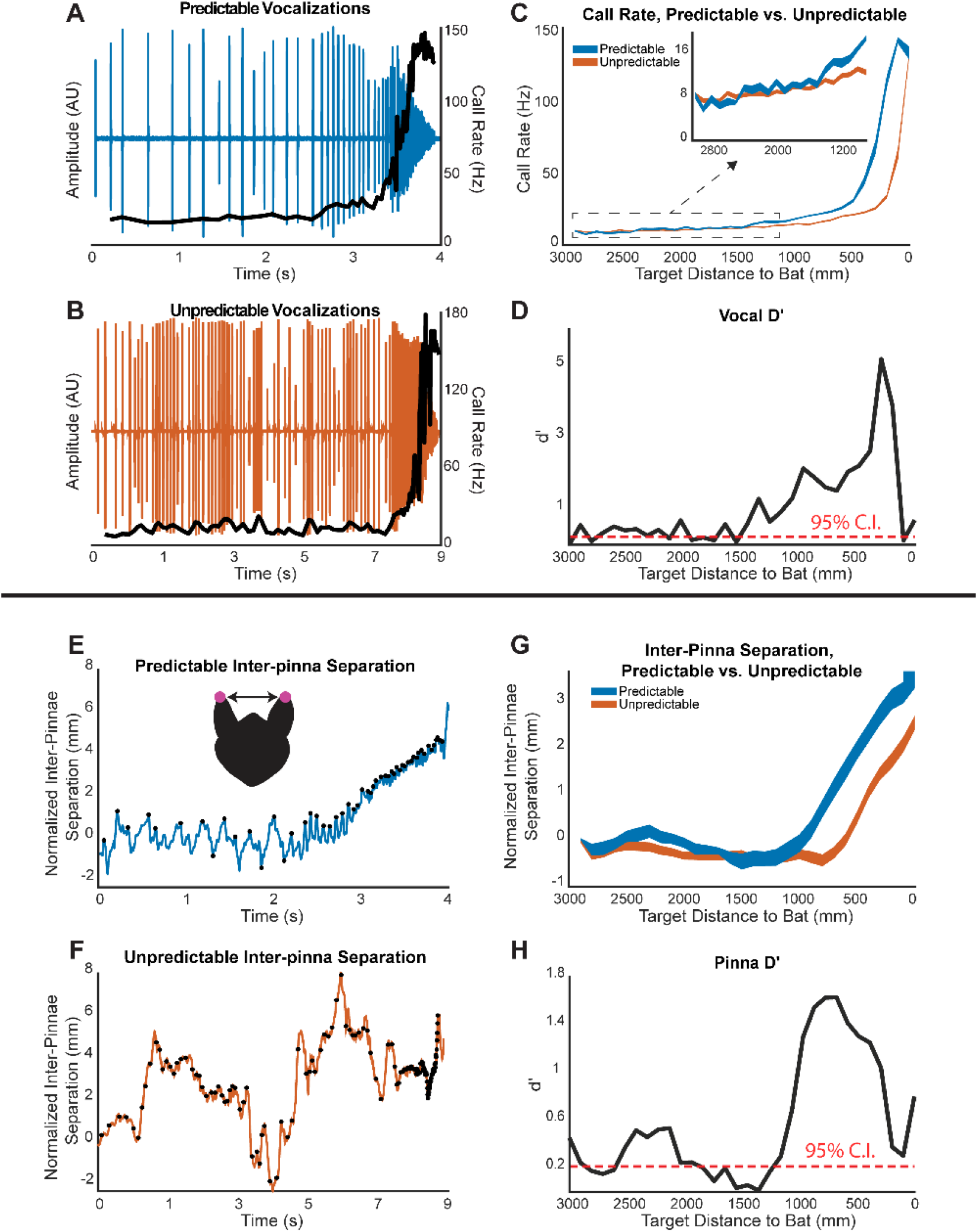
A-D) Vocal analysis. A) Example oscillogram of single trial data from a predictable motion trial. Call rate increases as the target approaches the bat as shown by black line superimposed over the oscillogram. B) Example oscillogram of single trial data from an unpredictable motion trial. Call rate increases as the target approaches the bat as shown by black line superimposed over the oscillogram. Note the different scale of the time axis compared to panel A. C) Mean call rate +/− standard error of the mean (s.e.m) plotted against target distance for predictable (blue) and unpredictable (orange) motion. D) d’ analysis of panel C with red dashed line indicating the 95% confidence interval (regions above are significant). E-H) Inter-pinna separation analysis. E) Example plot of normalized inter-pinna separation versus time for a single trial of predictable motion. Black dots denote times of echo arrival to indicate when the bat rapidly moves its ears during signal reception. F) Example plot of normalized inter-pinna separation versus time for a single trial of unpredictable motion (black dots indicate echo arrival times). Note the different scale of the time axis compared to panel E. G) Mean +/− s.e.m. change in inter-pinna separation for each motion condition. H) d’ analysis of panel G, with a red dashed line indicating the 95% confidence interval (regions above are significant).

Bats also adapt the positions of the pinna (the outer ear) based on changing target distance (Wohlgemuth et al., 2016). At further target distances, the ears are in an upright orientation with a narrow separation (see illustration in Figure 2E). As the target approached, the pinna flatten and reach a maximum inter-pinna separation at target arrival. Figure 2E shows the increase in inter-pinna separation as a function of decreasing target distance for a single predictable trial, and one unpredictable trial in 2F. In both panels, black dots indicate times of echo arrival. Prior work (Wohlgemuth et al, 2016) has shown that the bats perform these movements for each arriving echo to dynamically alter how the echo hits the outer ear. These movements represent short-term changes in positioning of the outer ears.

To compare how much the separation between pinna increases across trials and bats, we normalized the change in inter-pinna separation by calculating the average separation during the first 0.5 seconds of target motion and then subtracting this value from the raw values collected during the trial (see Methods). Using this method, all trials start at zero displacement and are then measured with respect to this baseline. The bats increase the inter-pinna separation as the target approaches, with the maximum change in separation being approximately 4 mm from the start of tracking to the end of capture. This represents an increase of approximately 20% in inter-pinna separation and is similar to values found in prior work (Wohlgemuth et al., 2016).

During the tracking phase, the bat moderately increases call rate with decreasing target distance, as shown in the expanded view in Figure 2C. The vocal behavior of the bat, in terms of call rate between the motion conditions, is not significantly different until the target is 1500 mm from the bat (d’ analysis in Figure 2C). The significant difference in call rate between predictable and unpredictable motions then continues through to capture. Importantly, we also observe similar differences in motor behavior with respect to inter-pinna separation control.

When comparing inter-pinna separation for predictable and unpredictable motion (Figure 2E-G) the bats maintain a relatively similar inter-pinna separation between predictable and unpredictable motion until the target distance is approximately 1200 mm from the bat, at this point the inter-pinna separation deviates significantly across the two motion types and stays different through target capture (Figure 2G and H). The data in Figure 2 show how target motion unpredictability delays the increase in inter-pinna separation and call rate that typically occurs at close target distances (~1200 mm for pinnae, ~1500mm for call rate). We also performed an ROC analysis on the data in Figure 2C and 2G to determine if the effects we found were dependent on the analysis approach. We find similar results for ROC and d-prime analyses: inter-pinna separation deviates at a target distance of ~1200 mm between predictable unpredictable motions. However, ROC analysis found that call-rates deviate at target distances closer to 2500 mm between the two motion conditions, 1 meter further than the d-prime analysis. Regardless, both analysis approaches identified significant differences in the adaptive control of call rate and inter-pinna separation as a function of target motion conditions.

The above results suggest unpredictable target motions delay increases in call rate and inter-pinna separation. Prior work shows that the transition from tracking phase to capture phase behaviors involves an abrupt increase in both call rate and inter-pinna separation (Wohlgemuth et al., 2016), and the above data suggest that this transition is delayed when tracking unpredictable motion. Delayed changes in behavior based on accumulated information are likely related to top-down differences in information processing, motivating an examination of adaptive behavioral controls related to changes in top-down sensory signaling. Sonar sound groups (SSGs) are bursts of calls produced at a higher rate and have been shown in past work to be associated with top-down influences on motor control (Kothari et al., 2018). On a behavioral level, prior work shows that bats increase the production of SSGs when encountering more challenging acoustic tasks, including erratically moving prey (Hulgard & Ratcliffe, 2016; Kothari et al., 2014). Figure 3A shows an example of the changes in call interval as a function of target distance. The ‘sawtooth’ nature of the line is indicative of SSG production, with short call intervals surrounded by longer call intervals. Black and blue dots on the plot identify SSGs with blue dots indicating a triplet (three calls) and black dots being doublets (two calls). Because of their quick repetition compared to the relatively longer call intervals of other calls surrounding them, SSGs during hunting are thought to facilitate rapid collection of information as the bat seeks to identify and localize their prey in preparation for capture (Kothari et al., 2014).

**Figure 3.**
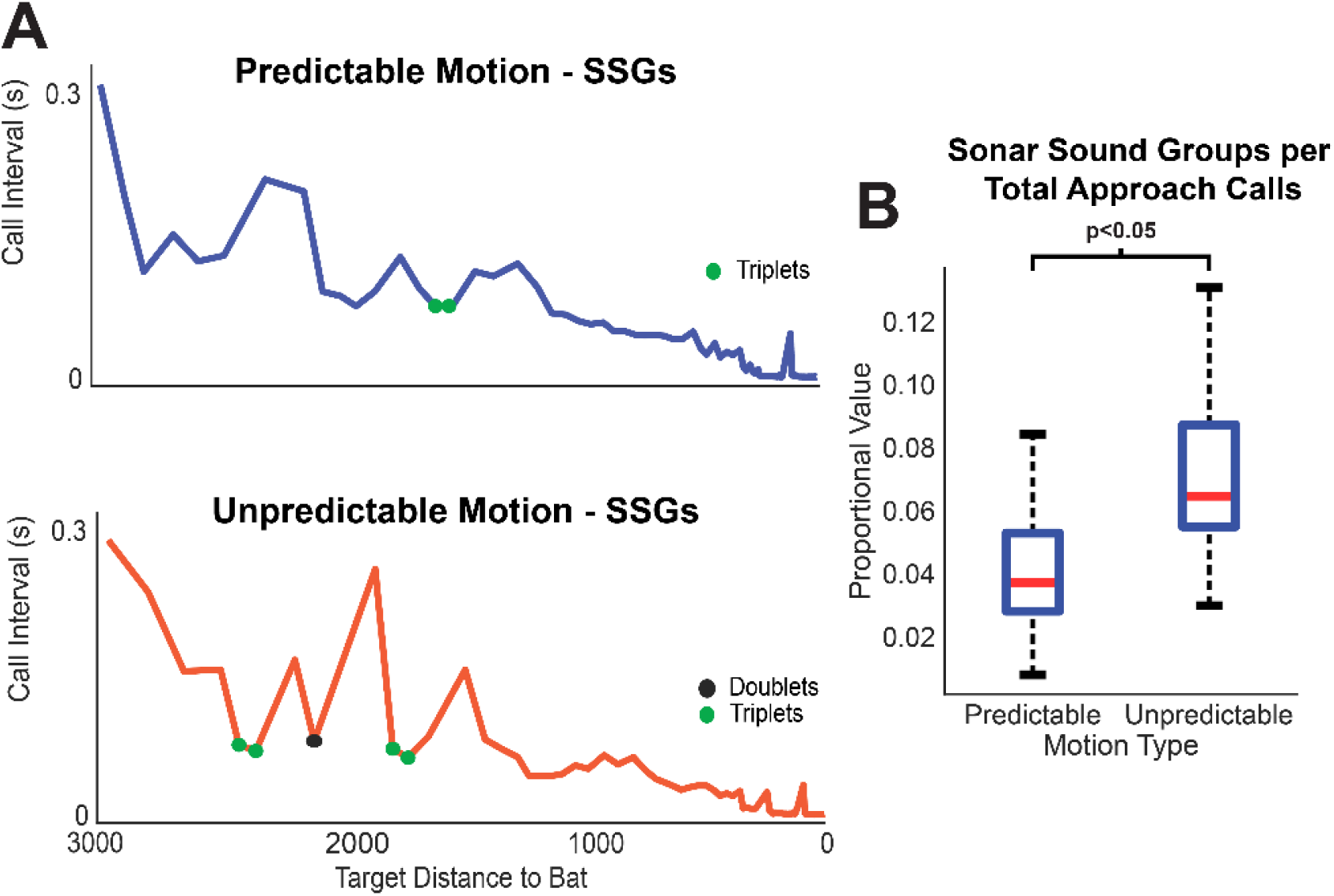
A) Top, call interval plotted as a function of target distance to the bat for a predictable motion trial. Colored dots indicate sonar sound groups or SSGs. Bottom, call interval plot for an unpredictable motion trial. B) Boxplot showing the usage of SSGs per the total number of calls for the tracking phase of the vocal pattern in each of the motion types. Unpredictable motion elicited significantly more SSGs than predictable motion (p<0.05, permutation test).

In past work, the proportion of calls in SSGs significantly increased when tracking erratically moving prey (Kothari et al., 2014). We performed the same SSG analysis on the predictable and unpredictable sonar call data when bats were in the tracking phase of the task in the current study (see examples of each trial type in Figure 3A). For this analysis, we look at the overall proportional change in SSG vs nonSSG production, not at the number of SSGs produced in sequence. Figure 3B shows a box plot comparing the proportion of calls included in SSGs for each trial type. The unpredictable motion trials are shown to have significantly (p<0.05, permutation test) more SSGs in the tracking phase compared to the predictable motion type, even though the overall call rate remained similar (Figure 2B). The median value for predictable motion was 4% SSG’s with a range from 1% to 9%. Unpredictable motion had a median value of 5% SSG’s with a range from 3% to 13%. Verified by a permutation test, the bat’s vocal-motor behavior during unpredictable motion elicited significantly more SSGs than during predictable motion, indicating a top-down influence on behavioral control related to target motion unpredictability. Interestingly, the production of SSGs for unpredictable motions was higher at the distances where the target changed direction (1-2 meters) than at the same distances for predictable motion (Figure S3). These results demonstrate how the bat increases SSG production when sensory uncertainty increases.

Past work has demonstrated top-down signals associated with the times of SSG production (Kothari et al, 2018). Considering the possibility of top-down influences driving SSG production, we examined whether changes in top-down signaling during the tracking phase have extended effects beyond the production of SSGs. Prior experiences influence top-down signaling (Sohoglu et al., 2012), and bats learn many aspects of their hunting strategies through interactions with their mothers and other conspecifics (Patriquin & Ratcliffe, 2023). Further, developmental work on bats found that young bats go after easier, low-quality prey early in life, and then with experience learn to hunt the more difficult but higher reward prey items (Aldasoro et al., 2024). We posit that properly timing the transition between hunting modes (e.g. tracking to capture phases) is experience dependent, and therefore under top-down control. More specifically, during the tracking phase the bat collects and updates information about the location of the target. This information is then used to appropriately time the transition from tracking to capture behaviors. This behavioral transition involves a sequence of behaviors, including an abrupt increase in call rate and pinna separation, as well as changes in flight and body positioning (Geberl et al., 2015; Wohlgemuth et al., 2016). We hypothesize that unpredictability in target motion during the tracking phase will delay the transition to the capture phase behaviors. By delaying this behavioral transition, the bat collects more information about the target’s position before initiating the capture sequence and presumably increases hunting success.

As mentioned earlier, the separation of the pinnae and call rate differ across predictable and unpredictable target motion conditions. For both motion varieties, the bat abruptly increases call rate and inter-pinna separation during the capture phase as seen in the example plot in Figures 2A and 2D, but there are differences in the timing of these behavioral changes across motion conditions. Analysis of capture-related pinna movements was conducted to identify the start of these capture behaviors with respect to target distance. Figure 4 shows examples of single trial vocal rate changes, plotted as a function of trial time, and inter-pinna separation over time for predictable (blue) and unpredictable (orange) motion trials. These data are plotted with respect to time to ease in visual comparison across motion conditions. The unpredictable moving target passes through the same target distances 3 times whereas the predictable only once, creating overlap in the unpredictable plots. Because the target’s motion is under experimental control, it is straightforward to convert trial time into distance in Figure 5. The green dot indicates the calculated start of capture-related behaviors for the respective motion type, and the dashed line shows the time of target arrival. This time, and corresponding target distance, can then be used to compare hunting mode transitions between target motion types and motor behaviors.

**Figure 4.**
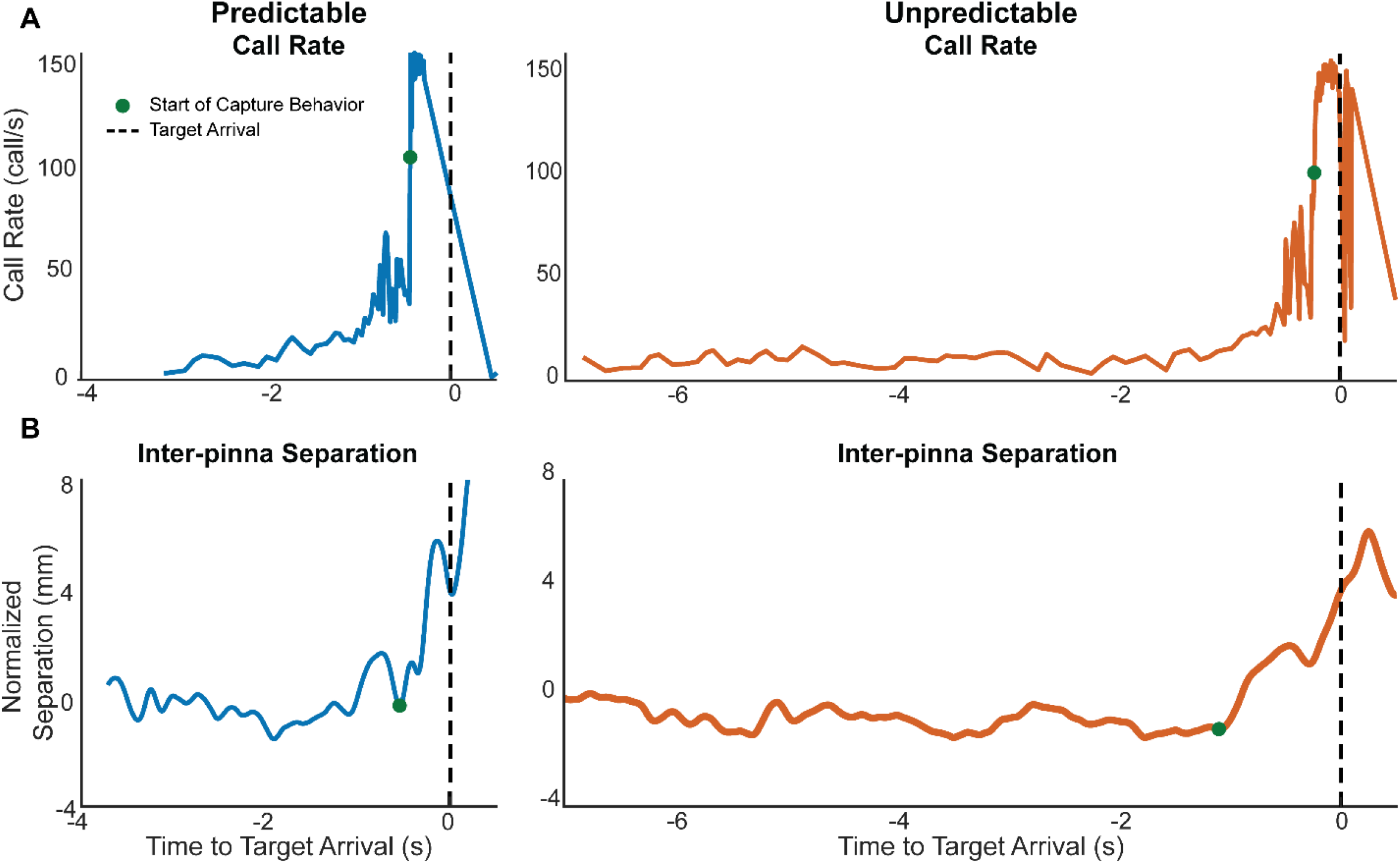
A) Example of call rate versus time for a predictable motion trial (left, plotted in blue), and an unpredictable motion trial (right, plotted in orange). The green dots indicate the calculated start of the capture phase, and the dashed black line indicates the time of target arrival. B) Example of smoothed inter-pinna separation versus time for predictable and unpredictable motion trials, plotted in blue (left) and orange (right), respectively. The green dot indicates the calculated start of the outward pinna movement, and the dashed black line indicates target arrival.

**Figure 5.**
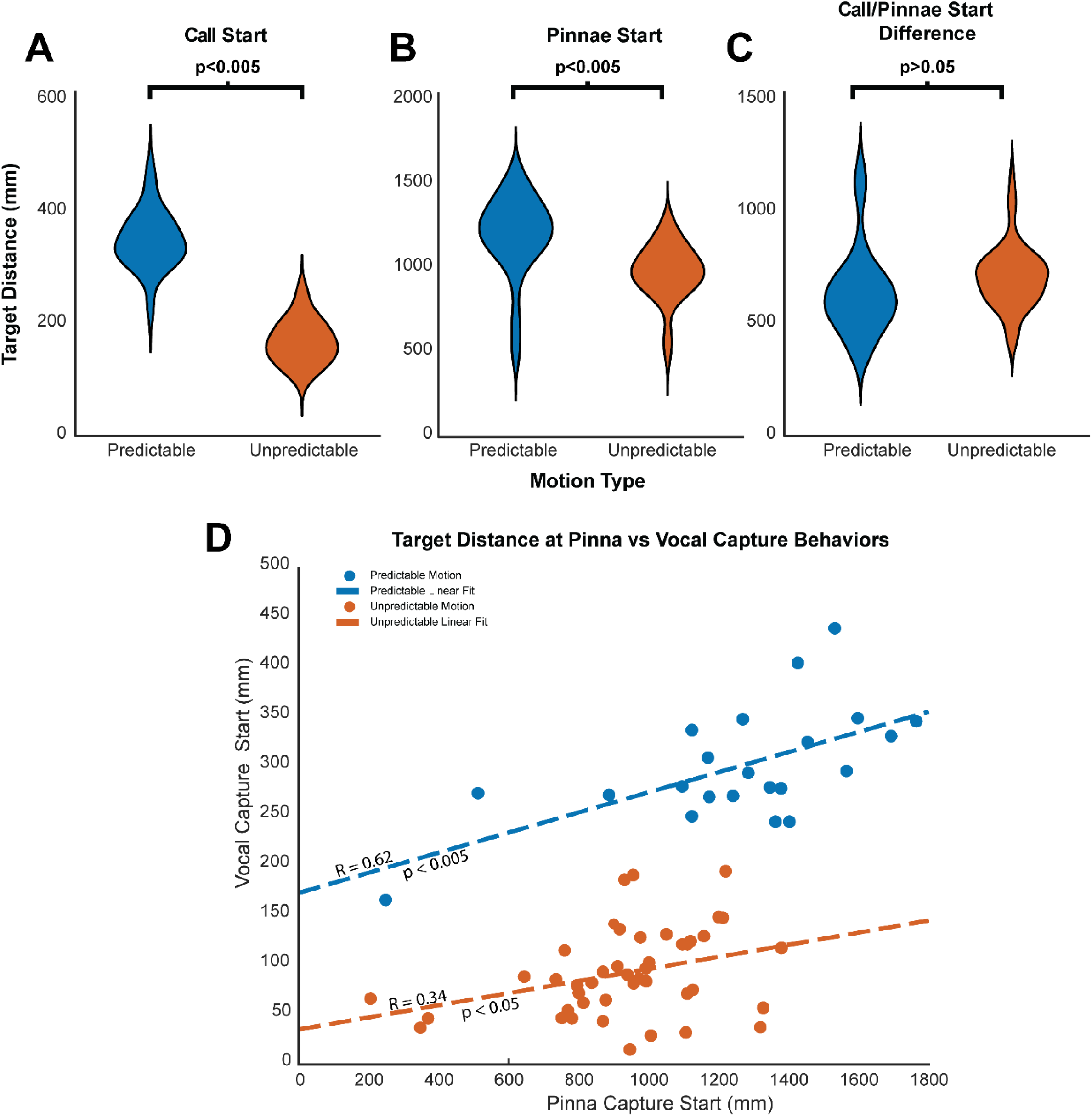
A) Violin plot comparing the target distances at which the bat transitioned its sonar call pattern into the capture phase. Unpredictable motion elicits a transition into capture at a significantly shorter distance than predictable (p<0.005, permutation test). B) Violin plot comparing the target distances at which the bat started widening its pinnae in preparation for target capture. The start of this movement begins significantly closer to the bat in unpredictable motion trials compared to the predictable motion trials (p<0.005, permutation test). C) Violin plot showing the difference in target distance from the start of the capture-related pinnae movements to the start of the capture-related call production. There is not a significant difference between the two motion types (p>0.05, permutation test). D) Scatterplot with the distance the pinnae begin their outward movement to start the capture phase plotted against the distances the call capture phase begins. Linearity is plotted in a color corresponding to the representative group and Pearson correlation coefficients are shown.

We used the methods outlined above to analyze the capture-related transitions in call rate and inter-pinna separation. Figures 5A and B summarize this analysis, showing that the bats transition into sonar call capture behaviors at shorter target distances for unpredictable motion trials as compared to predictable motion trials (i.e., they transition to capture phase behaviors *earlier and at further target distances* for predictable motion trials; p<0.001, permutation test, Figure 5A). The median target distance for the transition to capture phase call behaviors is 300 mm from the bat in predictable motion trials and 75 mm for unpredictable motion trials. There is little to no overlap in the range of distances where, excluding outliers, the closest target distance for predictable motion was 240 mm and the greatest for unpredictable motion was 190 mm. These data demonstrate a significant change in capture phase behaviors as a result of target motion predictability. Comparing these data to field and laboratory studies of bats hunting on the wing show similar results, but it is not possible to directly compare our study to past studies because these studies did not have location information for both the target and the bat. Field studies show a time duration of 200-500 milliseconds for capture phase, and in the laboratory similar time durations are found (Moss and Surlykke, 2001; Surlykke and Moss, 2000). Extrapolating from these times, and the 2-6 meter/second (Falk et al., 2015) flight speed of bats, would suggest that the transition to capture behaviors occurs less than 1 meter from the target. Our data are within this range.

There is a similar and significant effect of target motion unpredictability on capture phase behaviors for inter-pinna separation: inter-pinna movements related to capture were delayed for unpredictable motion as compared to predictable target motion (p<0.005, Figure 5B). For predictable motion trials, the median target distance at which inter-pinna separation increased in preparation for capture was 1300 mm, whereas this transition occurred at 900 mm for unpredictable motion trials. As is seen for adaptations to call rate, unpredictable target motion significantly delays the transition from tracking behaviors to capture phase behaviors for the control of pinnae positioning.

We also wanted to examine any correlated changes in call rate and inter-pinna separation. Within the same trial, the timing of the outward movement of the pinnae is tied to the start of the vocal capture phase. This can be characterized by looking at the difference in distance for the start of capture-related call and pinnae control. The difference in target distance between the start of the pinna movements and the start of the vocal capture phase is shown as a violin plot in Figure 5C, grouped by target motion type. There is not a significant difference between the two motion types (p>0.05, permutation test) relating to the timing of the pinnae with respect to the start of capture, demonstrating how these two behavioral events are tied to one another even if target motions vary. In other words, as the start of capture-phase behaviors in pinnae control shifts to closer distances, so too does the control of capture-related call behaviors.

As shown in Figure 5C, the difference in distances at which vocal and pinna-related capture behaviors begin is not significantly different between predictable and unpredictable motion. Figure 5D is a scatter plot with the distances at which the pinnae begin their capture-related movements plotted against the corresponding start distances of vocal capture behaviors. Linear fits are plotted in a color corresponding to the representative group and correlation coefficients are shown. There is a positive, significant correlation among predictable motion trials (R=0.62, p < 0.05), and a lower correlation, but still significant correlation, among unpredictable motion trials (R=0.34, p < 0.05). In order to test whether there were any outlier data points that could be affecting these results, we removed all data points more than 3 absolute median deviations away from the median. This identified one data point in the predictable motion condition (the left most blue data point at a target distance ~200 mm for inter-pinna separation). Upon removing this data point, the correlation coefficient was reduced, but still significant (r = 0.44, p < 0.05). These data suggest a common control mechanism mediating the transition to capture behaviors for pinnae motion and call production.

## Discussion

Three bats were trained on the ‘platform target tracking task’ (see Methods and Wohlgemuth et al, 2016 for a complete description) where the movement of the target can be experimentally controlled. Using this paradigm, the predictability of target motion was manipulated. We find the target motion predictability affects the bat’s adaptive sonar behaviors on both short and longer time scales. On a short time scale, we saw delayed changes in call rate and ear positioning for unpredictable motion as compared to predictable target motion. We also replicated earlier findings showing that bats increase the production of sonar sound groups (SSGs, a rapid burst of calls surrounded by calls at a lower rate) when tracking an erratic (i.e., unpredictable) target (Kothari et al, 2014). Additionally, we hypothesized that target motion predictability would influence behavioral control over longer timescale. Specifically, we predicted that the transition between the tracking phase and the capture phase of hunting would be delayed for unpredictable target motions. We find that both the abrupt increase in call rate, and the widening of inter-pinna separation related to capture preparation, are delayed for unpredictable target motions compared to predictable target motions (i.e., the changes happen at shorter distances for unpredictable target motions).

The data above demonstrate how target motion predictability influences behavior across two timescales: moment-by-moment changes leading to changes in both call rate and inter-pinna separation (e.g., delays in call rate and inter-pinna separation increases for unpredictable motion), as well as longer-term changes related to delays in the transition to capture behaviors (e.g., shorter distances for call rate and inter-pinna separation for unpredictable target motion). Importantly, the change in target motion predictability occurred at target distances greater than the distance at which bats transition from tracking to capture behaviors. This effect demonstrates how target motion predictability continues to influence behavior for up to several meters/seconds after the event. Additionally, we also found that unpredictable target motions increased the production of SSGs in a manner similar to prior work on bats as they tracked erratically moving prey (Kothari et al., 2014). These behavioral adaptations are presumably made to account for changes in target motion predictability and increase the probability of capture success.

This experiment also provides insight into the control of call rate increases with respect to the timing of inter-pinna separation increases. Overall, the dramatic increase in inter-pinna separation related to capture occurs at a target distance greater than the increase in call rate associated with capture behaviors (Figure 5A-C). However, this effect does not detail whether changes in target motion predictability first affect call rate or inter-pinna separation. As seen in Figure 2, call rate deviated across motion conditions at a target distance of ~1.5 meters, while inter-pinna separation deviated at ~1.2 meters (Figure 2E, d-prime significance crossing points). This result demonstrates how the effects of target motion predictability first alter call rate control, and then inter-pinna separation, suggesting that information collected from returning echoes first influences sonar call production and then influences pinna positions. Additionally, Figure 5D shows that the timing of call rate increases for the capture phase are correlated with the timing of inter-pinna separation for capture (although, the timing is reversed in this situation), further suggesting that these two behaviors are interconnected. Future research would benefit from focusing on disambiguating the temporal aspects of returning sonar echo influences on pinnae positioning and call rate adjustments.

One aspect of the current research that is important to consider is how target motion predictability influences the bat’s ability to anticipate the future location of the target. Importantly, all trial types were randomly interleaved, meaning that target motion predictability occurs within a trial and not across trials. For the predictable trials, the target moves at a consistent velocity and direction towards the bat for the entire target path. Past work in bats tracking targets along the horizontal axis shows that bats build an internal model of prey motion to aid in capture success (Salles et al., 2020). This work shows that bats can predict the movement of a target when it goes behind an acoustic occluder, and we posit that the bats in the current study are also creating internal models of the prey’s motion for predictions. For predictable motion trials, successively returning echoes provide feedback for a model of the prey approaching the bat. For unpredictable motion trials, however, the target unexpectedly changes direction with respect to the bat several times, resulting in a conflict between echo information and an internal model of *approaching* prey motion. What happens when a bat collects sensory information about the prey’s motion that conflicts with the bat’s internal model of prey motion? We hypothesize that in these circumstances, the bat must collect more information about the target’s position before preparing for prey capture. Past work on motor control in humans has found a similar ‘suppression’ of motor plans in response to ‘unexpected events’ (Wessel & Aron, 2017). The authors predict that for both action-dependent and action-independent (as is the case for the current study), a fronto-basal ganglia network produces a global signal that has a suppressive effect on motor output. We posit a similar top-down network delays the transition from tracking to capture behaviors. In doing so, the bat collects more information about the location of the target to increase hunting success.

We examined the average target distance at which the bat transitions from target tracking to target capture behaviors for predictable and unpredictable moving prey. We defined this transition as the abrupt increase in both call rate and inter-pinna separation right before target arrival. We developed an algorithm that identified when the bat reached its maximum call rate and then back-calculated the start of that increase in call rate (see Methods). We employed a similar approach to identify the beginning of the abrupt increase in inter-pinna separation related to the transition to capture behaviors. Although there can be some noise in using these methods to find the exact start of the capture phase, we applied the same approach to both motion conditions to calculate comparatively when capture behaviors began across motion varieties. Using this method, we find that capture behaviors are indeed delayed when tracking an unpredictably moving target (see Figures 4 and 5). This result lends support to the hypothesis that bats must collect more sensory information from an unpredictably moving target before transitioning to capture behaviors, resulting in a capture transition at closer target distances than when tracking a predictably moving target.

Our research identified short-term changes in sonar behaviors related to target motion predictability (i.e., call rate, inter-pinna separation), as well as effects that extended over significantly longer timeframes (i.e., the transition from tracking to capture behaviors). Considering this outcome, we suggest that the short-term changes in behavior are driven by bottom-up networks, most likely inputs from the inferior colliculus to the superior colliculus. In this schema, the inferior colliculus processes auditory cues related to the target’s position and then passes this information to the superior colliculus for instantaneous sensorimotor control. There are also obvious influences of top-down circuits in the bats’ behaviors. Recruiting top-down networks would help the bat in using predictions as well as more dynamic motor adaptations to a changing sensory landscape. Similar to past work, we find an increase in SSG production when tracking unpredictably moving targets (Kothari et al., 2014). Prior work on the superior colliculus shows an increase in the gamma power of the local field potential during SSG production (Kothari et al, 2018), a signal generated by cortical, top-down inputs (Canolty et al., 2006). Therefore, we suggest that unpredictable target motion increases SSG production as well as top-down signaling from the cortex to the midbrain. We argue that this top-down signal initiated by SSG production is ultimately what delays the transition from tracking to capture behaviors. In this way, manipulations of target motion predictability allow us to disambiguate parallel processing for short- and long-term sensorimotor control.

One additional aspect of this study that should be discussed is that the bat is not flying towards the prey item in this experiment, the prey item is flying towards the bat. This condition may alter the way in which the bat is adapting its behavior as compared to flying bats. A foraging bat flies typically flies at 4-6 meters per second (Falk et al., 2014); whereas the typical moth flies much slower, at 0.25 to 1.0 meters per second. As such, bats are often flying faster than their prey, and therefore closing the distance between themselves and the target. However, there are insects that have evolved to hear the echolocation calls of bats, and will perform evasive maneuvers when they hear the bat getting close to them (Ratcliffe et al., 2009). Interestingly, echolocating bats will produce more SSGs when tracking evasive prey (Kothari et al., 2014), a finding that is recapitulated in the current study. Moreover, past work using a similar stationary bat paradigm demonstrated changes in call rate and duration similar to those seen in flying bats hunting on the wing (Aytekin et al., 2010). We therefore conclude that the behavioral changes as a result of unpredictable prey motion in the current study are relevant to flying bats.

The goal of the current study was to characterize how animals change their behavioral strategies when tracking predictable versus unpredictable moving targets. We hypothesized that an animal must collect more information about an unpredictable moving target in preparation for target capture than a target that is moving along a predictable motion path. We find that echolocating bats do indeed delay their capture behaviors to collect more information when tracking an unpredictable moving target. Importantly, the bats also changed their behaviors on a moment-by-moment basis when tracking an unpredictable moving target, showing the dual timescale effects of unpredictability. Future work would benefit from focusing on underlying neural structures that may be contributing to the bottom-up and top-down influences on short-term and long-term sensorimotor control, respectively.

## Acknowledgments

We would like to thank Cynthia Moss, the Johns Hopkins animal facility staff, and undergraduates in the Moss Lab at Johns Hopkins University for helping with data collection.

## Data and resource availability

All relevant data and resources can be found within the article and its supplementary information

---

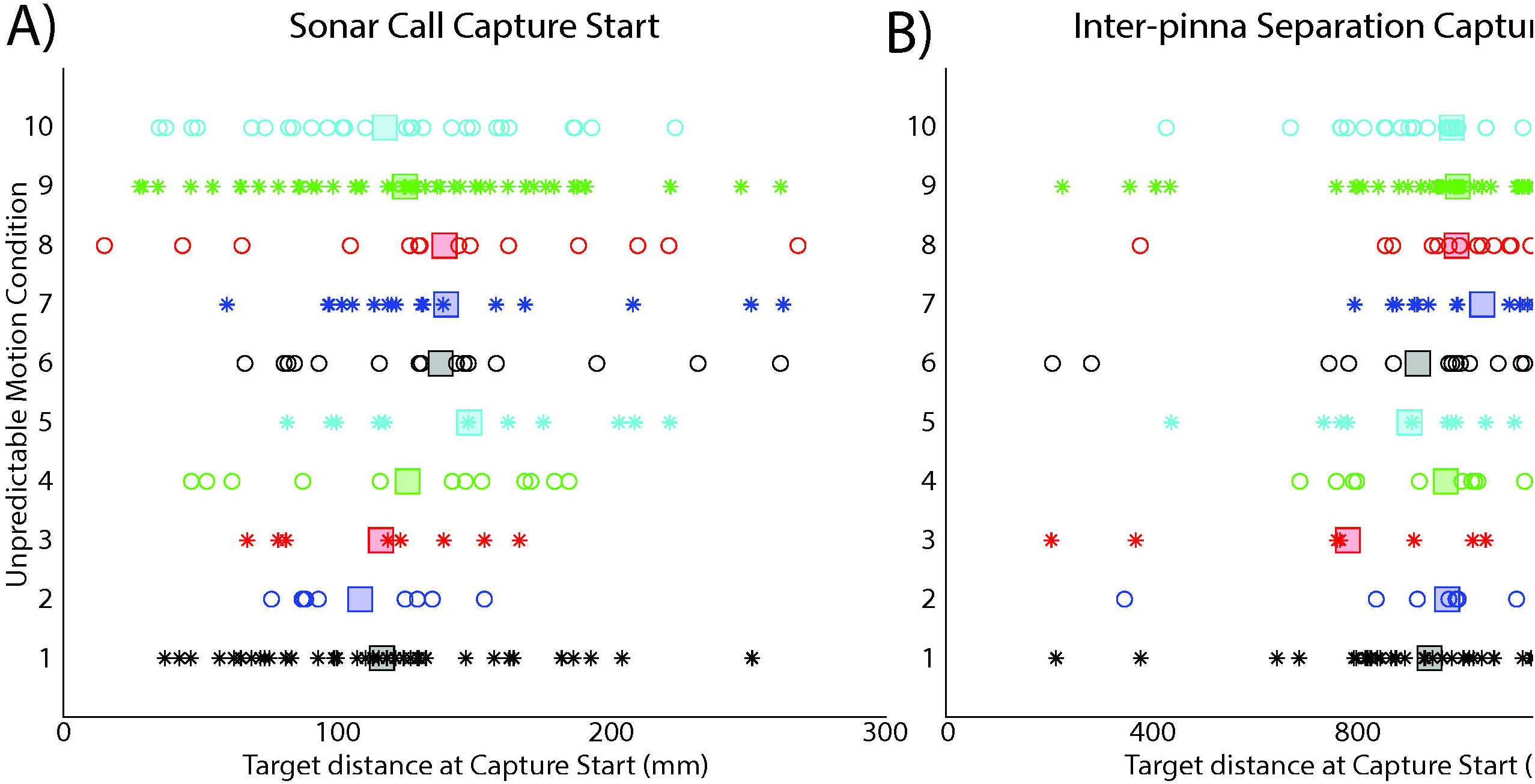

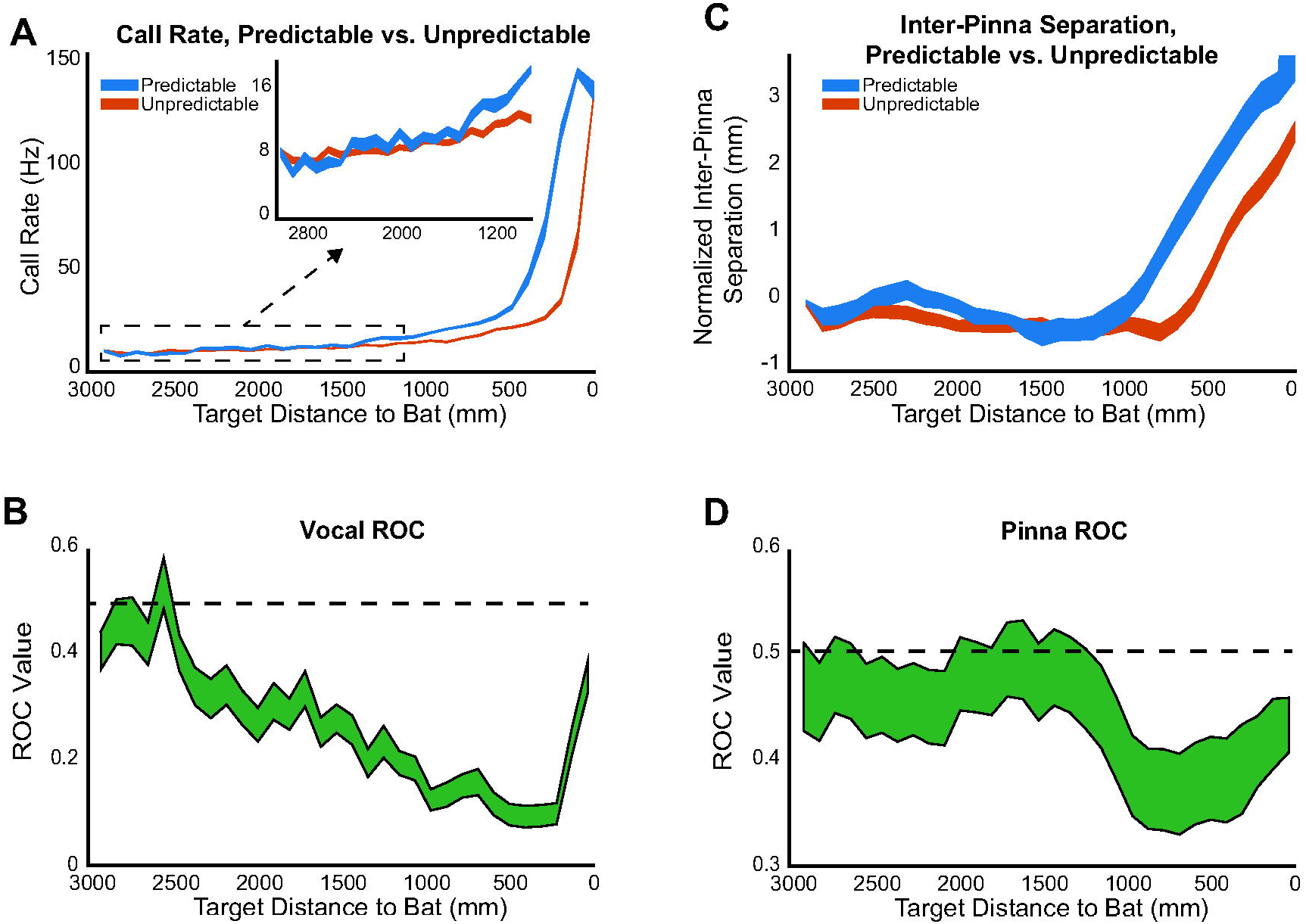

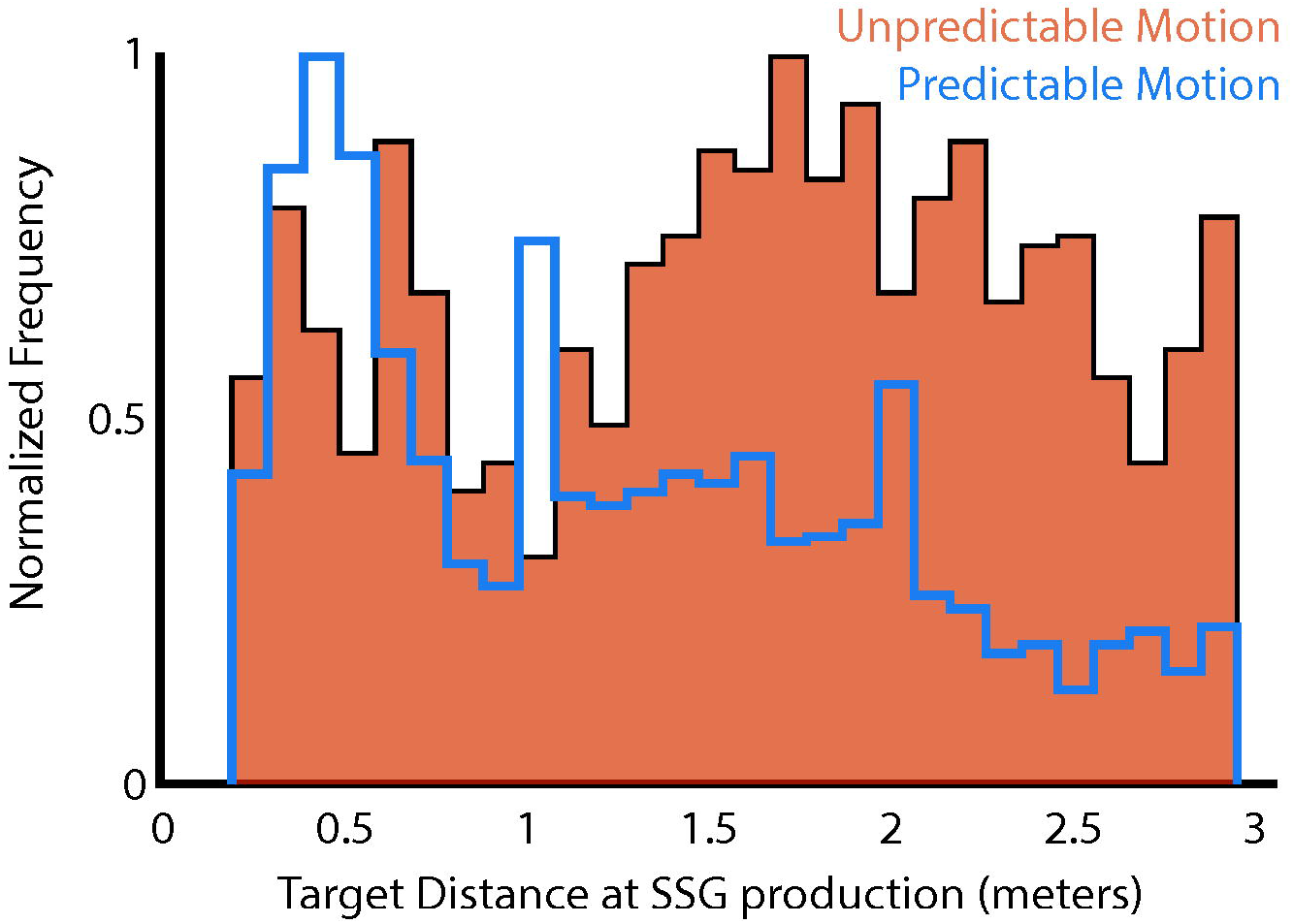

