## Supplemental Figures for "Unpredictable Prey Motion Shapes Behavior Across Timescales"

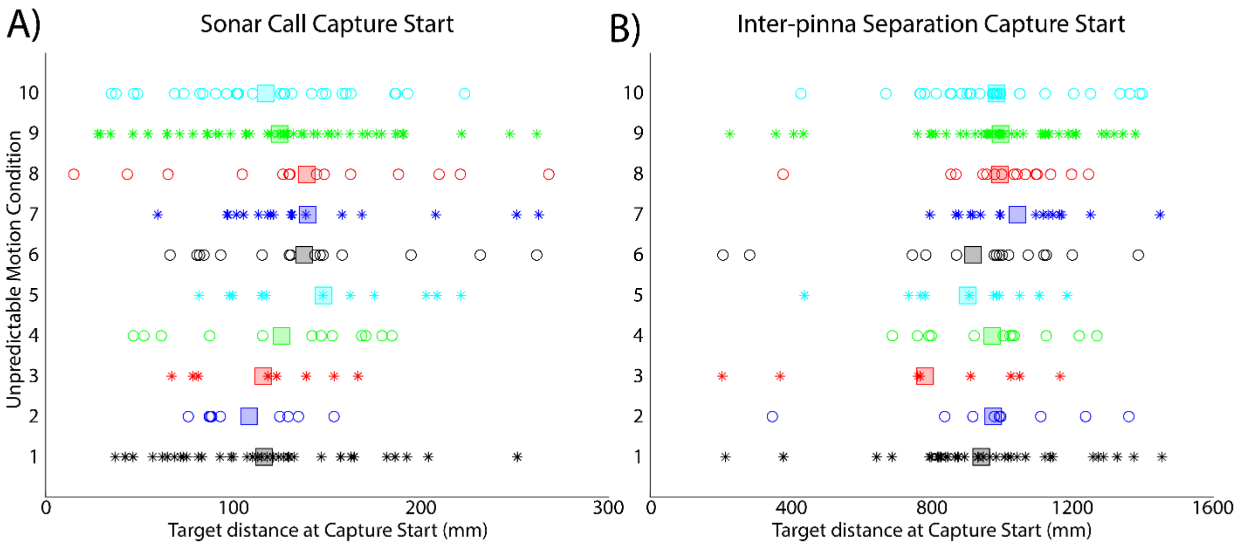


**Figure S1:** A) Target distances (millimeters) at which the bat transitioned from tracking to capture behaviors for sonar call control across all unpredictable motion conditions (n = 202). The average target distance for each motion condition is plotted as a large, shaded square. There are no significant differences in target distance for the transition to capture behaviors across unpredictable motion conditions (1-way, repeated measures ANOVA, $p > 0.05$). B). Target distances (millimeters) at which the bat transitioned from tracking to capture behaviors for inter-pinna separation across all unpredictable motion conditions (n = 202). The average target distance for each motion condition is plotted as a shaded square. There are no significant differences in target distance for the transition to capture behaviors across unpredictable motion conditions (1-way, repeated measures ANOVA, $p = 0.34$).

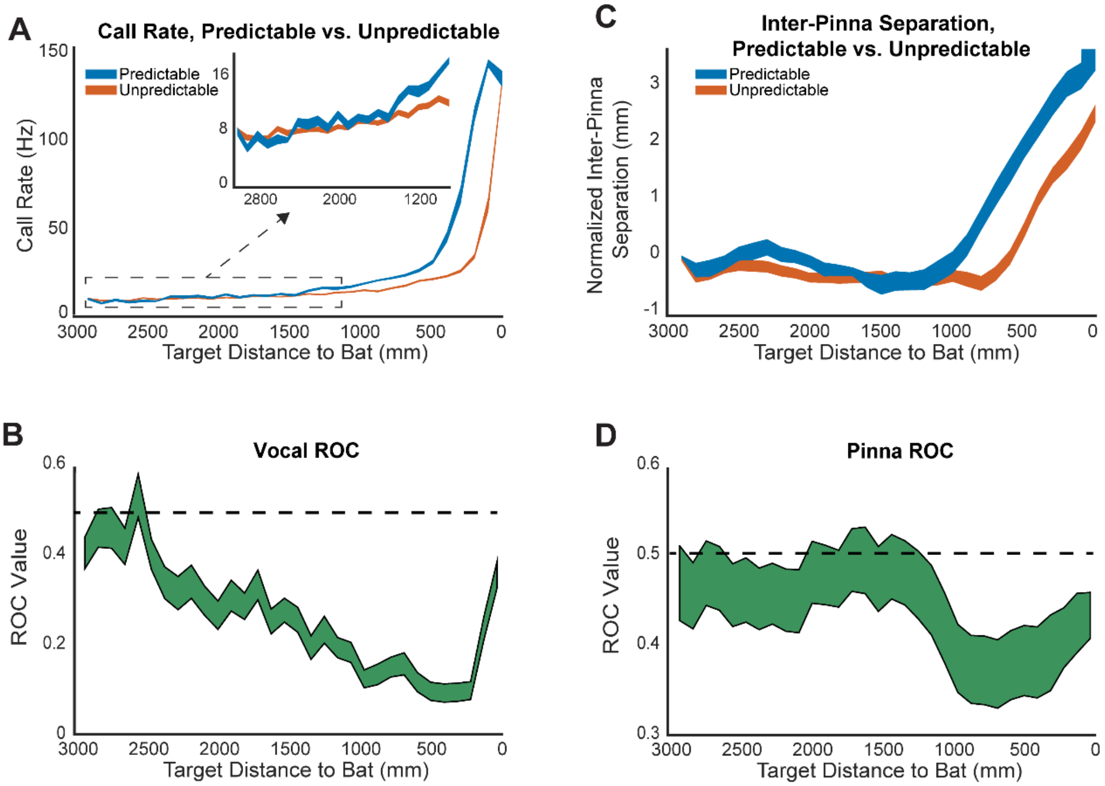


Figure S2: A) Replotting Figure 2C showing the change in call rate as a function of target distances for predictable (blue) and unpredictable (orange) motions. B). ROC analysis performed on data from Figure S2A binned in 10-centimeter bins. Plotted is the mean ROC value +/- the standard error for each bin. C) Replotting Figure 2C showing the change in interpinna separation as a function of target distances for predictable (blue) and unpredictable (orange) motions. D) ROC analysis performed on data from Figure S2C binned in 10-centimeter bins. Plotted is the mean ROC value +/- the standard error for each bin.

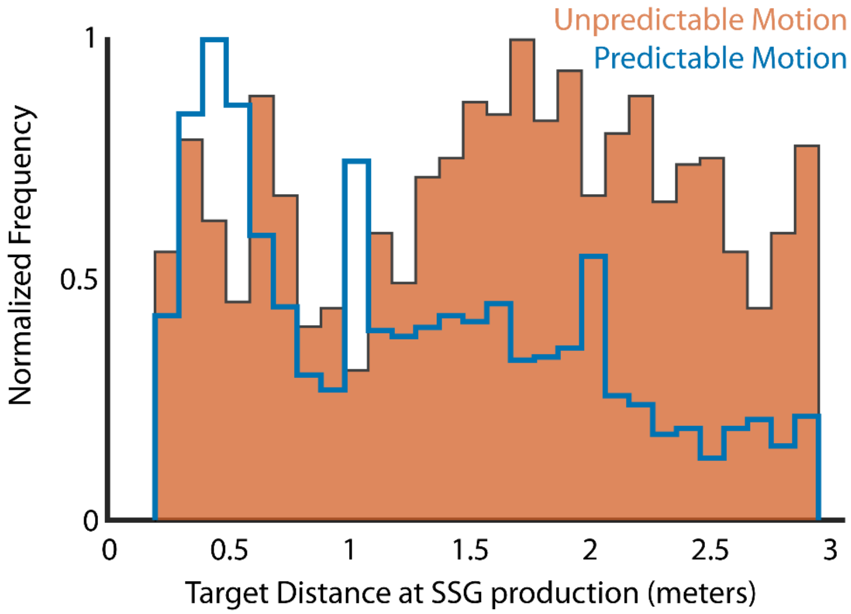


**Figure S3:** Histogram showing the target distances at which SSGs were produced for predictable motion (blue) and unpredictable motion (orange). The mean distance at which SSGs were produced for unpredictable motion was significantly greater than predictable motion (permutation test, $p < 0.05$).
